# Ligand-Mediated Biofilm Formation via Enhanced Physical Interaction Between a Diguanylate Cyclase and Its Receptor

**DOI:** 10.1101/212183

**Authors:** David Giacalone, T. Jarrod Smith, Alan J. Collins, Holger Sondermann, Lori J. Koziol, George A. O’Toole

## Abstract

The second messenger, cyclic dimeric GMP (c-di-GMP) regulates biofilm formation for many bacteria. The binding of c-d¡-GMP by the inner-membrane protein LapD controls biofilm formation, and the LapD receptor is central to a complex network of c-di-GMP-mediated biofilm formation. In this study, we examine how c-di-GMP signaling specificity by a diguanylate cyclase (DGC), GcbC, is achieved via interactions with the LapD receptor and by small ligand sensing via GcbC’s <u>cal</u>cium channel <u>che</u>motaxis (CACHE) domain. We provide evidence that biofilm formation is stimulated by the environmentally relevant organic acid citrate (and a related compound, isocitrate) in a GcbC-dependent manner through enhanced GcbC-LapD interaction, which results in increased LapA localization to the cell surface. Furthermore, GcbC shows little ability to synthesize c-di-GMP in isolation. However, when LapD is present GcbC activity is significantly enhanced ~8-fold, indicating that engaging the LapD receptor stimulates the activity of this DGC; citrate-enhanced GcbC-LapD interaction further stimulates c-di-GMP synthesis. We propose that the I-site of GcbC serves two roles beyond allosteric control of this enzyme: promoting GcbC-LapD interaction and stabilizing the active conformation of GcbC in the GcbC-LapD complex. Finally, given that LapD can interact with a dozen different DGCs of *P. fluorescens*, many of which have ligand-binding domains, the ligand-mediated enhanced signaling via LapD-GcbC interaction described here is likely a conserved mechanism of signaling in this network. Consistent with this idea, we identify a second example of ligand-mediated enhancement of DGC-LapD interaction that promotes biofilm formation.

**Importance:** In many bacteria, dozens of enzymes produce the dinucleotide signal c-di-GMP, however it is unclear how undesired crosstalk is mitigated in the context of this soluble signal, and how c-di-GMP signaling is regulated by environmental inputs. We demonstrate that GcbC, a DGC, shows little ability to synthesize c-d¡-GMP in the absence of its cognate receptor LapD; GcbC-LapD interaction enhances c-di-GMP synthesis by GcbC, likely mediated by the I-site of GcbC. We further show evidence for a ligand-mediated mechanism of signaling specificity via increased physical interaction of a DGC with its cognate receptor. We envision a scenario wherein a “cloud” of weakly active DGCs can increase their activity by specific interaction with their receptor in response to appropriate environmental signals, concomitantly boosting c-di-GMP production, ligand-specific signaling and biofilm formation.

## Introduction

For most bacteria, biofilm formation is a highly regulated event (1, 2). The bacterial intracellular second messenger, cyclic dimeric GMP (c-di-GMP) controls biofilm formation by regulating a diversity of biofilm-relevant outputs (3), including flagellar motility (4), extracellular polysaccharide production (5, 6), adhesin localization (7), and transcriptional control of pathways important for early biofilm formation (8). An important research theme in the field has been understanding the mechan¡sm(s) of c-di-GMP signaling-specificity in the context of microbes that can have >50 proteins that make, degrade and bind this second messenger.

Biofilm formation by *Pseudomonas fluorescens* Pf0-1 occurs when the adhesin LapA localizes to the cell surface (9). LapA is maintained on the cell surface when the inner membrane protein LapD binds c-di-GMP. The c-di-GMP-bound, inner membrane-localized LapD sequesters LapG, thus this protease is unable to target the N-terminal cleavage site of LapA (10, 11). One example of how c-d¡-GMP is specifically transferred to the LapD receptor is by physical interaction with a diguanylate cyclase (DGC) (12). The DGC called GcbC has been shown to physically interact with LapD utilizing a surface exposed α-helix of the GGDEF domain on GcbC and a surface exposed α-helix of the EAL domain of LapD (12). We also demonstrated previously that the I-site of GcbC contributes to the interaction of this enzyme with LapD (13). This direct interaction model was proposed as one means to confer signaling specificity (12, 14).

GcbC is an inner membrane protein that contains a putative <u>cal</u>cium channel <u>che</u>motaxis receptor (CACHE) domain N-terminal to its GGDEF domain; this CACHE domain is predicted to reside in the periplasm. CACHE domains can be responsible for small ligand sensing (15). Many signal transduction proteins, including DGCs and histidine kinases, contain CACHE domains, which are involved in modulating these enzyme activities (16,17,18). GcbC, along with five other DGCs encoded on the *P. fluorescens* Pf0-1 genome (Pfl01_1336, Pfl01_2295, Pfl01_2297, Pfl01_3550, and Pfl01_3800), contain putative periplasmic CACHE domains located N-terminally to their GGDEF domain, suggesting that these six DGCs are capable of sensing and responding to small ligands. The predicted domain organization of three of these CACHE-containing proteins, GcbC, Pfl01_2295, and Pfl01_2297, is shown in Figure 1A.

**Fig. 1.**
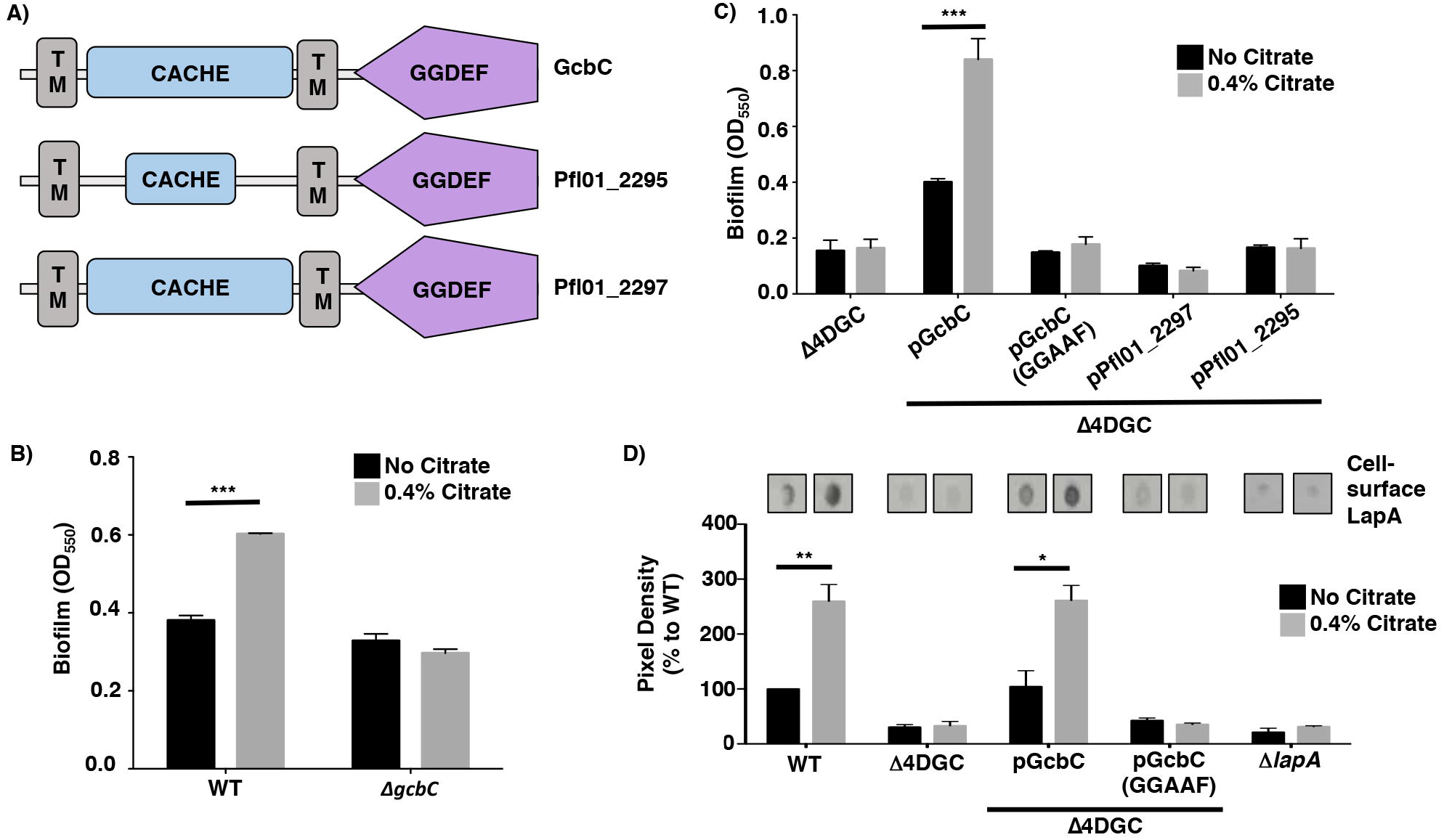
Citrate stimulates biofilm formation via GcbC. (A) Predicted domain organization of GcbC, Pfl01_2295, and Pfl01_2297 as predicted by SMART (34). The domains are indicated in each block. TM = transmembrane domain. Analysis of Pfl01_2295 by MiST2 (35) predicts a second transmembrane domain. (B) Biofilm formation by WT *P. fluorescens* and the *ΔgcbC* mutant in the presence and absence of 0.4% citrate (n = 3; + SD). (C) Biofilm formation by the indicated strains +/-citrate. In this panel, the Δ4DGC mutant background is used, with the WT GcbC and the catalytically inactive variant (GGAAF) introduced into this mutant on plasmids. Pfl01_2297 and Pfl01_2295 are two other CACHE-domain containing diguanylate cyclases in *P. fluorescens*, expressed from the same vector backbone, that serve as controls. (D) Quantification of cell-surface levels of LapA in the presence and absence of 0.4% citrate. Representative blots are shown. For panels B, C, and D, the values shown are an average of 3 replicates (+ SD); horizontal black bars indicate a P value of <0.05 (*), <0.01 (**), or <0.001 (***) with a student’s t-test compared to the condition without citrate.

In this study, we analyze the putative CACHE domain of GcbC and identify two environmentally relevant organic acids, including citrate, as a ligand for this DGC. We show citrate-enhanced physical interaction of GcbC and LapD, and that this enhanced interaction promotes increased c-di-GMP synthesis by GcbC, thereby promoting biofilm formation. We also describe another example of such ligand-mediated enhancement of biofilm formation, suggesting the generality of this mechanism. Furthermore, we show that GcbC has little propensity to synthesize c-di-GMP in isolation; it is only in the presence of its receptor, LapD, that GcbC makes high levels of c-di-GMP. We propose that the I-site of GcbC serves two roles beyond allosteric control of this enzyme: promoting LapD-GcbC interaction and stabilizing the active conformation of GcbC in this complex. Thus, we propose a mechanism whereby GcbC engagement with its receptor LapD, which is enhanced by environmental cues, could serve as a general mechanism to confer specificity to this complex signaling network.

## Results

### Citrate-Mediated Biofilm Enhancement is Dependent on GcbC Activity

Automated domain annotation software predicts a CACHE domain in the periplasmic portion of GcbC (Figure 1A). To identify small molecules that the CACHE domain of GcbC may bind, we performed a CLUSTAL alignment of this amino acid sequence with CACHE domains of known structure (19). GcbC showed the highest amino acid similarity to rpHKlS-Z16 (PDB ID: 3LIF) of *Rhodopseudomonas palustris* with 31% identity (Fig. S1). When crystallized, the CACHE domain of rpHKlS-Z16 was bound to citrate and methyl-2,4-pentanediol, two small ligands recruited from the crystallization cocktail (19). Because ligand binding by CACHE domains has been shown to activate the C-term¡nally fused histidine kinase (HK) and DGC domains (17, 18), we hypothesized citrate may stimulate GcbC activity to enhance biofilm formation.

To test this idea, we compared the impact of citrate on biofilm formation in the wild type *P. fluorescens* Pf0-1 and a *gcbC* mutant. In a wild-type (WT) background, a 50% increase in biofilm formation was observed in the presence of citrate (Fig. 1B). However, citrate-mediated enhancement of biofilm formation observed for the wild-type (WT) strain was abolished in the *gcbC* mutant (Fig. 1B), suggesting that citrate-mediated enhancement of biofilm formation is dependent on the presence of GcbC. Also, we found that increasing concentrations of citrate further enhanced biofilm formation of *P. fluorescens* by up to ~100%. (Fig. S2A)

Next, we selected the CACHE domain-containing Pfl01_2295 and Pfl01_2297 proteins (Fig. 1A), both of which interact with LapD (20), to serve as controls to determine if other CACHE domains also respond to citrate to promote biofilm formation. For these studies, we used a *P. fluorescens* Pf0-1 strain lacking four DGCs, referred to as Δ4DGC, which does not form a biofilm under our laboratory conditions; this low c-di-GMP producing strain was previously used to investigate c-di-GMP production by other *P. fluorescens* DGCs (21). The four DGCs deleted in the Δ4DGC strain, identified previously, are *gcbA, gcbB, gcbC*, and *wspR* (21). Of the three CACHE domain-containing DGCs expressed in the Δ4DGC mutant background (GcbC, Pfl01_2295, and Pfl01_2297), only the strain expressing GcbC responded to citrate with increased biofilm formation (Fig. 1C). DGC-mediated c-di-GMP synthesis typically requires an intact GGDEF active-site motif and mutation of this motif to GGAAF in GcbC eliminates catalytic activity; this GcbC (GGAAF) mutant variant was shown previously to be stably expressed at a level equivalent to the WT protein (21). When GcbC-GGAAF mutant protein was expressed in the Δ4DGC mutant background, biofilm formation was abolished and notably, citrate-mediated biofilm formation was also abolished (Fig. 1C).

We further assessed the functionality of GcbC, Pfl01_2295, and Pfl01_2297 as active DGCs via assaying their ability to stimulate increased production of the Pel polysaccharide using congo red (CR) binding assay. Increased CR binding indicates increased production of the Pel polysaccharide, which is mediated by increased c-di-GMP production, thus dark red colonies indicate robust Pel production and white colonies indicate a lack of Pel production. Each DGC was cloned into an expression plasmid, then electroporated into *P. aeruginosa* PA14 and grown under inducing conditions with 0.1% arabinose. When GcbC and Pfl01_2295 are expressed from a plasmid, CR binding of Pel polysaccharide was robustly stimulated compared to the vector only control (pMQ72, Fig. S3). When Pfl01_2297 was expressed from a plasmid, CR binding of Pel polysaccharide was, at best, weakly stimulated compared to the vector only control. As a control, a mutant defective in production of the Pel polysaccharide *(ΔpelA)* showed little CR binding (Fig. S3). These data indicate that Pfl01_2295 and Pfl01_2297 are likely functional DGCs, a point addressed further below.

Importantly, citrate-enhanced biofilm formation is dependent on LapA (Fig. S2B), indicating that citrate acts via the known LapD-LapG-LapA pathway. To further support a role for LapA in citrate-stimulated biofilm formation, we used *P. fluorescens* Pf0-1 strains carrying a HA-tagged LapA variant to detect the amount of LapA at the cell surface, as reported (7, 10), as a function of the presence of citrate. In a WT *P. fluorescens* strain, citrate caused a 159% increase in LapA pixel density, which suggests a higher abundance of LapA at the cell surface in the presence of citrate (Fig. 1D). Only when GcbC was present and catalytically active did citrate cause an increase in cell surface-associated LapA (Fig. 1D). As a control, minimal levels of signal were detected in a *P. fluorescens* strain lacking LapA, as expected. Also, there was no detectable difference of the amount of LapA in whole cell lysate in the presence and absence of citrate in WT *P. fluorescens* (Fig. S2C). Taken together, these data show that citrate-mediated stimulation of biofilm formation by WT *P. fluorescens* Pf0-1 requires the active diguanylate cyclase GcbC and is associated with enhanced cell-surface LapA.

Citrate also enhances growth of *P. fluorescens* (Fig. S2D), but as described below, this enhanced growth is not the basis for the enhanced biofilm formation in the presence of citrate. Interestingly, despite its ability to stimulate biofilm formation, citrate caused a ~100% increase in swimming motility (Fig. S2E), perhaps via the ability of citrate to enhance growth of *P. fluorescens.*

### The Putative Ligand-Binding Site of the CACHE Domain is Important for GcbC-Mediated Biofilm Formation

CACHE domains are ubiquitous, periplasmic, ligand-binding domains (15, 16). In a previous study, the RXYF motif was found to be the most conserved feature among the characterized CACHE domains (19). A mutation of the RXYF motif of the CACHE domain of KinD, a histidine kinase in *Bacillus subtilis*, caused this microbe to lose its ability to respond to root exudates, and resulted in decreased biofilm formation on tomato roots compared to a WT strain (17).

We first assessed whether citrate-enhanced biofilm formation was due to increased protein expression level. We measured protein levels of a HA-tagged GcbC variant (expressed from a plasmid from the studies shown in Figures 1C and 1D) in the presence and absence of citrate and found that citrate did not affect the production and/or stability of GcbC (Fig. 2A). Analysis of the pixel density of the bands +/-citrate revealed a ratio of 1.05+0.08, indicating no change in GcbC protein level when the cells are grown with citrate.

**Fig. 2.**
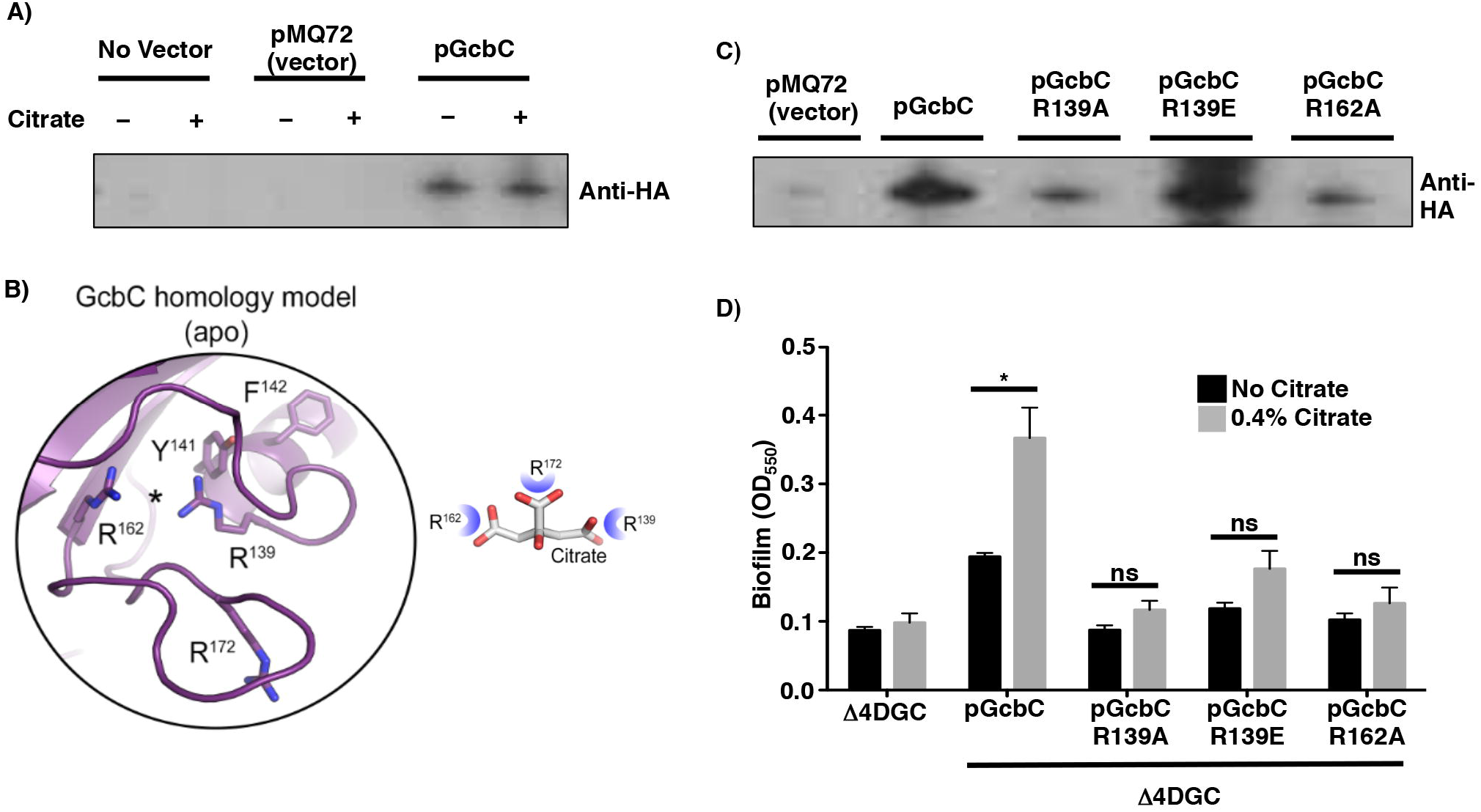
Effects of CACHE domain mutations on biofilm formation. (A) Western blot to assess the level of GcbC-HA expressed from a plasmid in the presence and absence of citrate. This is the GcbC-HA-expressing plasmid used in Figure 1. (B) Homology model of the CACHE domain of GcbC based on rpHK1S-Z16 (PDB ID: 3LIF) and vpHK1S-Z8 (PDB ID: 3LID) (see Figure S1 and S4) to identify the putative ligand-binding site. (C) Western blot assessment of the relative stability of GcbC-R139A-HA, GcbC-R139E-HA and GcbC-R162A-HA mutant proteins compared to the WT GcbC. These proteins were all expressed in Δ4DGC mutant *P. fluorescens* strain. (D) Quantitative analysis of biofilm formation by strains expressing the indicated CACHE domain mutants in the presence and absence of citrate. For panels A and E, data shown is the average of three replicates (+ SD); horizontal black bars indicate a P value of either <0.05 (*) or <0.001 (***) with a student’s t-test comparing each strain without citrate versus with added citrate, ns, not significant.

We next sought to identify the putative site where citrate might bind to GcbC using the known structures of CACHE domains (Fig. 2B, left; template PDB ID 3LIB). Like in the other CACHE domains, the tyrosine residue of the RXYF motif of GcbC was predicted to point towards the ligand-binding site (Fig. 2B, Fig. S4). Furthermore, based on the CACHE domain model of GcbC, the amino acids R139, R162, and R172 were identified as candidates to shape the predicted ligand-binding site (Fig. 2B, right). Based on the model, we predict that three arginine residues can coordinate the three carboxylic acid groups of citrate (Fig. 2B, right). R139 is also part of the RXYF motif. To probe whether these residues were involved in the citrate-mediated enhancement of biofilm formation, each of the arginine residues forming the putative ligand-binding site was mutated in GcbC and the mutant protein expressed from a plasmid introduced into the Δ4DGC mutant background. Mutating R172 (R172A and R172E) resulted in loss of stability of GcbC (Table S1), however GcbC-R139A, GcbC-R139E, and GcbC-R162A variants were detected by Western blot, with GcbC-R139E variant present at approximately WT levels (Fig. 2C).

We next assessed whether the CACHE domain was required for citrate-enhanced biofilm formation. The strain carrying GcbC-R139E variant, as well as the less stable GcbC-R139A and GcbC-R162A alleles, did not show a significant enhancement of biofilm formation in the presence of citrate (Fig. 2D). We further expanded our search for important conserved residues within the CACHE domain that were predicted based on alignments with other CACHE domain proteins, however the other twenty-seven mutant proteins we constructed were unstable (Table S1). Together, our data indicate that the RXYF motif, and specifically R139 of the putative citrate-binding arginine triad, is critical for GcbC-dependent, citrate-mediated enhancement of biofilm formation.

### Selected Organic Acids Stimulate Biofilm Formation via GcbC

We next sought to identify other ligands that may be sensed by the CACHE domain of GcbC, given that some characterized CACHE domains bind multiple ligands (19, 22). Based on the ability of the putative arginine triad to bind the three carboxyl groups of citrate, we tested whether other organic acids (Fig. S5A), including acetate (C2), pyruvate (C3), succinate (C4), fumarate (C4), α-ketoglutarate (C5) or isocitrate (C6) could enhance biofilm formation in a GcbC-dependent manner. In both a WT *P. fluorescens* strain and when GcbC is expressed in a Δ4DGC mutant strain, only isocitrate significantly enhanced biofilm formation, and did so by 40% (Fig. S5B). Like citrate, isocitrate is a C6 compound which contains three carboxyl groups (Fig. S5A). Also, isocitrate did not stimulate biofilm formation by the strain carrying the GcbC-R139E variant (Fig. S5C), indicating that the R139E variant was not responsive to citrate or isocitrate. Taken together, these data indicate that the CACHE domain of GcbC likely binds citrate and isocitrate.

### Citrate-Mediated Interaction of GcbC with LapD Enhances Synthesis of c-di-GMP

In our published model, GcbC mediates biofilm formation by transferring c-di-GMP to LapD through physical interaction, as demonstrated by both pull-down and bacterial two-hybrid assay (12). We further showed that the LapD-GcbC interaction is mediated via the α5^GGDEF^ helix of GcbC with the α2^EAL^ helix of LapD (12). This LapD-GcbC interaction also requires the I-site of GcbC (13). We asked whether citrate might exert its effect of stimulating biofilm formation via stabilization of the LapD-GcbC signaling complex. To test if citrate bolsters GcbC-LapD interaction, we exploited the bacterial two-hybrid system used to initially demonstrate interaction between these proteins (12). We did observe a modest but significant enhancement of LapD-GcbC interaction in the presence of citrate (Fig. 3A). No such enhancement was observed for the control interactions: LapD-Pfl01_2295, or GcbC with a dual domain protein (Pfl01_0192; this protein has cytoplasmic GGDEF and EAL domains similar to LapD) or a phosphodiesterase (Pfl01_2920, Fig. 3A). Citrate also did not enhance GcbC-GcbC dimerization (Fig. 3A). Further, the catalytically inactive variant of GcbC still showed citrate-stimulated interaction similar to WT GcbC (Fig. 3A). Finally, the GcbC-R139E mutant variant did not show a significant, citrate-mediated increased interaction (Fig. 3A), consistent with our data above that this residue is required for citrate-mediated enhancement of biofilm formation (Fig. 2D).

**Fig. 3.**
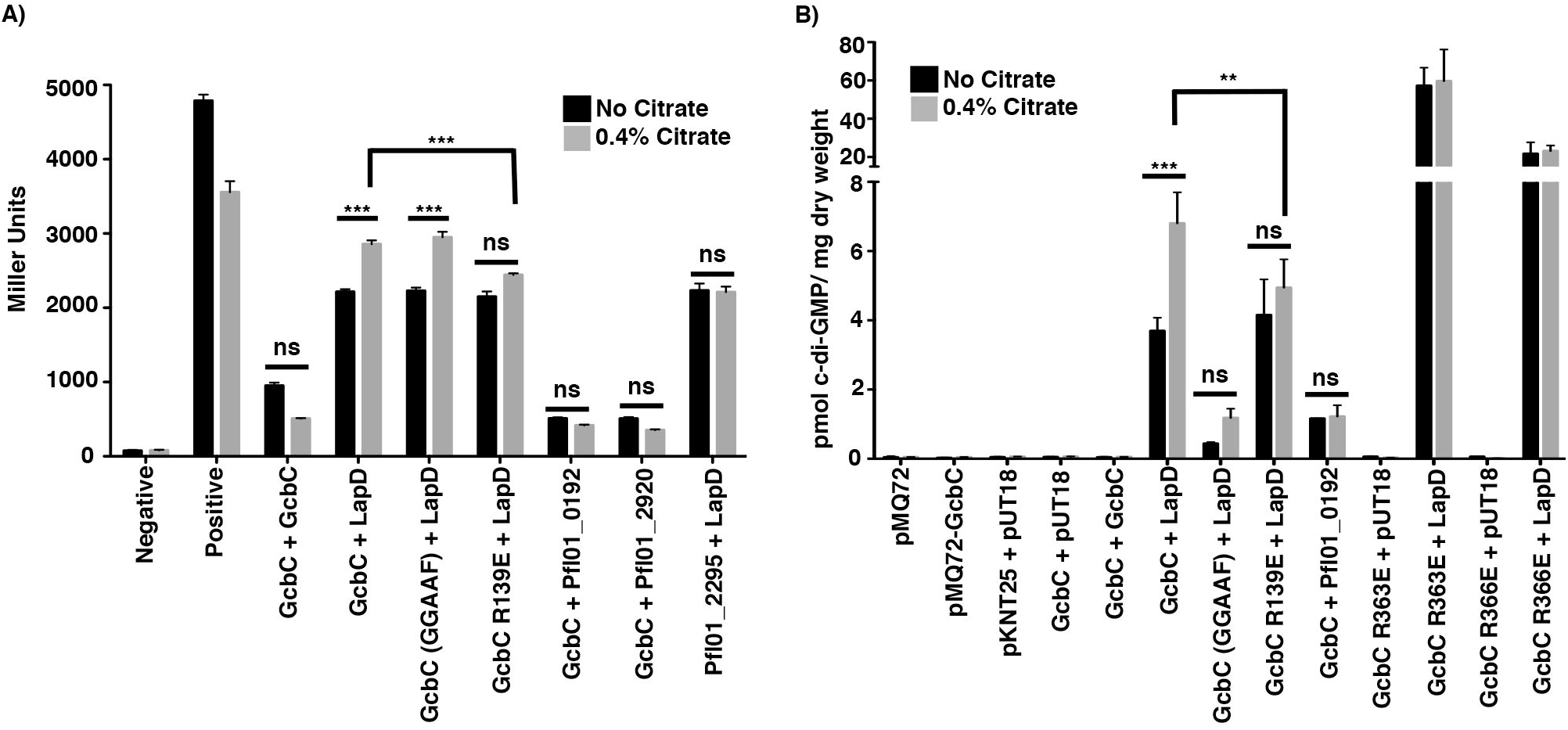
Citrate-mediated biofilm enhancement. (A) The effect of citrate on interaction with the indicated proteins is compared by the B2H assay, and expressed as Miller Units. Pfl01_2295 = CACHE domain-containing DGC; Pfl01_0192 = GGDEF and EAL domain-containing protein; Pfl01_2920 = EAL domain-containing protein. Briefly, the proteins of interest fused to two halves of the catalytic domain of an adenylate cyclase from *Bordetella pertussis* (T25 and T18) are transformed into *E. coli* BTH101 cells (33). Plasmids pKNT25 and pUT18 are the vector-only controls. If the two proteins interact, the adenylate cyclase activity is reconstituted, promoting cAMP synthesis, which in turn activates transcription of the *lacZ* gene and increases production of its gene product. The β-galactosidase activity is assessed using ortho-nitrophenyl-β-galactoside (ONPG) as a substrate, which serves as a measure of the degree of interaction (33). Citrate did not enhance interaction of the vectors or of the positive control (GCN4 leucine zipper protein). Citrate only enhanced the interaction between LapD and GcbC. The effect of catalytic activity of GcbC on interaction with LapD was compared in the presence and absence of 0.4% citrate. Mutation of the putative citrate-binding pocket in the CACHE domain of GcbC (GcbC-R139E) resulted in the loss of citrate-stimulated GcbC-LapD interaction. (B) The B2H assay strains were used to measure the effects of citrate on c-di-GMP synthesis by GcbC. Plasmids were cotransformed as indicated into *E. coli* BTH101. After 24 hours of incubation at 30°C, nucleotides were extracted. C-di-GMP production in *E. coli* BTH101 was determined for strains producing the indicated proteins. Only co-expression of LapD and GcbC showed appreciable accumulation of c-di-GMP, which was significantly stimulated by added citrate. For panels A and B, all experiments were performed in triplicate (+ SD), with an indicated P value of either <0.05 (*) or <0.01 (**) with a student’s t-test comparing each strain without versus with citrate, ns, not significant. For panel B, GcbC-LapD and GcbC R139E-LapD interactions are represented by six biological replicates instead of three.

We further explored whether citrate could also enhance the diguanylate cyclase activity of GcbC. We used the bacterial two-hybrid (B2H) plasmids and strains to express GcbC and LapD outside of their native context, and to better focus on how the interaction of these two proteins might specifically impact GcbC’s activity. The activity of GcbC was assessed by measuring the level of c-di-GMP extracted from the *E. coli* strains that carry plasmids containing the indicated genes (Fig. 3B). The level of c-di-GMP measured in the *E. coli* strain carrying the vector control was <0.5 pmol c-di-GMP/mg dry weight of bacteria (Fig. 3B). This low background of c-di-GMP provided a useful tool to measure differences in c-di-GMP levels derived from GcbC in the presence and absence of citrate and the LapD receptor.

GcbC alone did not synthesize levels of c-di-GMP above background and citrate did not promote c-di-GMP production using three different strains wherein only GcbC was expressed (Fig. 3B; pMQ72-GcbC, pKNT25-GcbC + pUT18, pKNT25-GcbC + pUT18-GcbC), a finding consistent with a previous study in which low levels of c-di-GMP synthesis by GcbC were detected (21). Co-expression of GcbC with LapD resulted in ~4 pmol c-di-GMP/mg dry weight, an ~8-fold increase above the control level, and the level of c-di-GMP was further, significantly increased to ~7 pmol c-di-GMP/mg dry weight upon addition of citrate (Fig. 3B). This finding has two important implications. First, GcbC activity is apparently stimulated when engaging its cognate LapD receptor, suggesting a mechanism for specific signaling between GcbC and LapD. Second, enhanced interaction of GcbC and LapD upon addition of citrate further enhances c-di-GMP production by GcbC.

Modest amounts of c-di-GMP catalysis occurred when the catalytically inactive GcbC variant was coexpressed with LapD, but the addition of citrate did not significantly enhance the level of c-di-GMP (Fig. 3B). Additionally, Pfl01_0192, a dual GGDEF-EAL domain protein with a cytoplasmic domain organization similar to LapD, which was shown to interact weakly with GcbC (Fig. 3A), produced levels of c-di-GMP in the presence of GcbC that were above background but did not increase with added citrate (Fig. 3B).

We investigated the impact of the GcbC-R139E CACHE-domain mutation on citrate-mediated LapD-GcbC interaction and c-di-GMP synthesis. While the GcbC-R139E/LapD interaction was equivalent to the interaction between LapD and the wild-type GcbC, citrate-enhanced interaction of GcbC-R139E-LapD showed a small, but non-significant increase (Fig. 3A). This finding was consistent with the data above (Fig. 2D), showing that the GcbC-R139E mutant was still capable of promoting biofilm formation in the Δ4DGC background, but biofilm formation was not enhanced by the addition of citrate. Furthermore, and consistent with the bacterial two-hybrid data, basal c-di-GMP levels were not affected by the GcbC-R139E mutant variant compared to the WT GcbC when coexpressed with LapD (Fig. 3B). Importantly, there was no significant increase in c-di-GMP level when the GcbC-R139E mutant variant was coexpressed with LapD in the presence of citrate (Fig. 3B). Thus, our data indicate that LapD-GcbC interaction enhances c-di-GMP production, and the addition of citrate stimulates both interaction of LapD-GcbC and c-di-GMP synthesis, likely via the CACHE domain of GcbC.

When GcbC is removed from its native context and expressed in a heterogenous system, this enzyme did not produce levels of c-di-GMP above background (Fig. 3B). This finding suggested that GcbC might be inactive until it engages its cognate receptor LapD. To test this idea, we assessed c-di-GMP production by I-site mutant variants of this enzyme expressed in the B2H *E. coli* strain. The canonical role of the I-site is to reduce GcbC catalytic activity when c-di-GMP is bound, and we previously identified two different mutants of GcbC in our near the I-site, R363E and R366E, which greatly enhanced biofilm formation and c-di-GMP levels when expressed in *P. fluorescens* (13). These I-site-proximal mutations also reduced interaction with LapD (13). When expressed on its own, the GcbC-R366E I-site mutant was not able to synthesize c-di-GMP levels above background (Fig. 3B). We observe a similar phenotype for the R363E mutant; R363 stabilizes the GGDEF domains as they come together in a GGDEF dimer with c-di-GMP at the I-site. Only when LapD was present were high levels of c-di-GMP produced, with GcbC-R363E-expressing strain synthesizing ~60pmol c-di-GMP/mg dry weight and GcbC-R366E-expressing strain synthesizing ~20pmol c-di-GMP/mg dry weight (Fig. 3B). These levels of c-di-GMP were significantly higher than the level detected for wild-type GcbC, and citrate did not further enhance c-di-GMP production for either I-site mutant (Fig. 3B). These data strongly support the model that GcbC has little or no activity until this enzyme engages its cognate receptor.

### A Conserved Mechanism of Signaling Specificity

We next explored if the mechanism we defined for GcbC-LapD interactions might apply to any of the other 20 DGCs in *P. fluorescens.* We found that LapD is a central hub of DGC interaction; LapD interacts with nine different DGCs in a pairwise bacterial two-hybrid assay (Fig. 4A,C). Included among the nine DGCs that interact with LapD are the CACHE domain-containing DGCs Pfl01_2295 and Pfl01_2297, and the putative SadC homolog, Pfl01_4451 (Fig. 4A). SadC is a DGC identified for its role in the early stages of biofilm formation by *P. aeruginosa* (23, 24), and the *ΔsadC* mutant of *P. aeruginosa* shows approximately a 50% reduction of global, cellular c-di-GMP levels compared to WT *P. aeruginosa* PA14 (24).

**Fig. 4.**
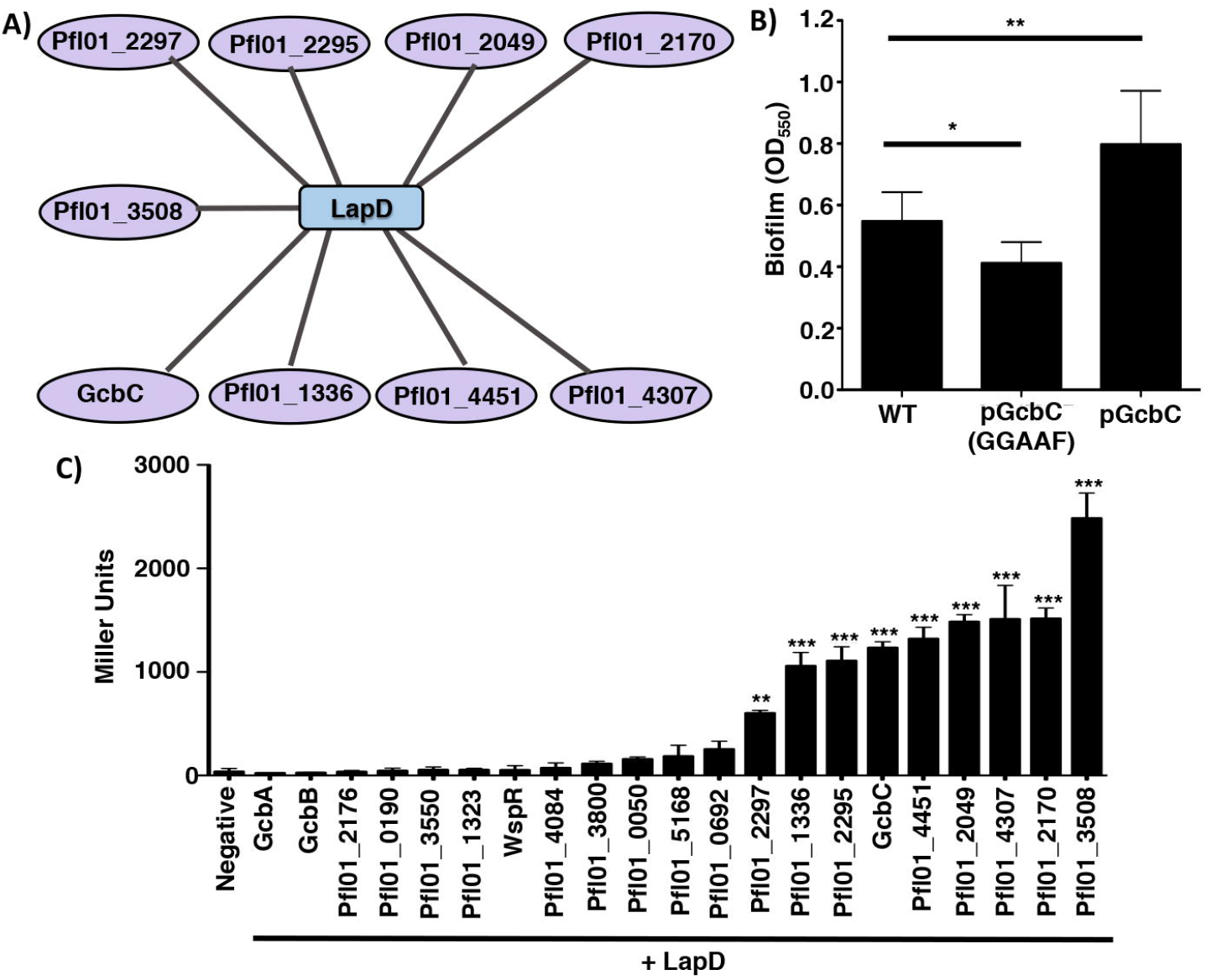
LapD interacts with multiple DGCs. (A) Interaction map of DGCs that showed a significant level of interaction with LapD by 24 hours, as compared to the negative control, by B2H assay. Data that served as the basis for this model is shown in Figure 4C. (B) Biofilm formation of WT *P. fluorescens.* WT GcbC and the catalytically inactive variant (GGAAF) were expressed on plasmids. Experiments were performed in triplicate (+ SD), with an indicated P value of either <0.05 (*) or <0.01 (**) with a student’s t-test comparing each strain to WT *P. fluorescens.* (C) Plasmids containing each of the 21 DGCs and LapD were cotransformed into *E. coli* BTH101. After 24 hours of incubation at 30°C, cells were scraped from the plate and β-galactosidase levels were measured to identify LapD interaction partners. Of the 21 DGCs tested, 9 were identified as significantly interacting with LapD by B2H assay compared to the vector only (negative) control. Pfl01_2297, Pfl01_1336, Pfl01_2295, GcbC, Pfl01_4451, Pfl01_2049, Pfl01_4307, Pfl01_2170, and Pfl01_3508 significantly interacted with LapD, but the level of interaction with LapD varied amongst the 9 DGCs. Assays were performed in triplicate with two biological replicates (+ SD). Significance was evaluated by a one-way ANOVA analysis followed by a Tukey Multiple Comparison Analysis test, comparing each DGC-LapD interaction to the negative control. Asterisks indicate a P value of either <0.01 (**) or <0.001 (***).

Given that there are nine DGCs that interact with LapD, we hypothesized that each of the nine DGCs may compete with one another to interact with LapD. To test this model, we overexpressed WT GcbC and the catalytically inactive variant of GcbC in a WT *P. fluorescens* strain. When the catalytically inactive GcbC variant was overexpressed, biofilm formation was significantly reduced by ~20% (Fig. 4B), indicating that the catalytically inactive variant of GcbC is displaying a dominant negative phenotype, a finding consistent with our competition hypothesis. As a control, when WT GcbC was overexpressed, biofilm formation was significantly enhanced, as expected (Fig. 4B).

We next explored if the ligand-mediated signaling specificity mechanism is conserved among other CACHE-containing DGCs. We focused on Pfl01_2295 given that this DGC shows similar levels of interaction with LapD as was observed for GcbC (Fig. 4C), and the likely robust c-di-GMP synthesis activity as indicated by the robust CR binding observed when Pfl01_2295 was expressed from a plasmid in *P. aeruginosa* (Fig. S3). Also, the CACHE domain of Pfl01_2295 contains the conserved RXYF motif (Fig. S6), and contains two of the three arginine residues that comprise the putative ligand binding site of the CACHE domain of GcbC. Thus, we predicted that Pfl01_2295 would sense ligands that are similar to, but different from, citrate. To test this idea, we expressed Pfl01_2295 in the Δ4DGC mutant background and performed a biofilm assay with medium supplemented with exogenous acetate, pyruvate, succinate, fumarate, or isocitrate. Citrate was shown not to stimulate biofilm formation through Pfl01_2295 nor enhance LapD – Pfl01_2295 interaction as measured using the bacterial two-hybrid assay (Fig. 1C, Fig. 3A).

Succinate significantly enhanced biofilm formation in the presence of Pfl01_2295 and also significantly enhanced interaction with LapD (Fig. 5A). Thus, this finding mirrored what we observed for citrate and isocitrate with GcbC. Interestingly, acetate significantly stimulated Pfl01_2295-dependent biofilm formation, but did not stimulate interaction with LapD (Fig. 5B), perhaps indicating that acetate stimulates Pfl01_2295 activity but not the ability of this DGC to interact with its receptor. Pyruvate, fumarate, and isocitrate had no effect in stimulating biofilm formation in a Δ4DGC mutant background with Pfl01_2295 expressed from a plasmid (Fig. S7).

**Fig. 5.**
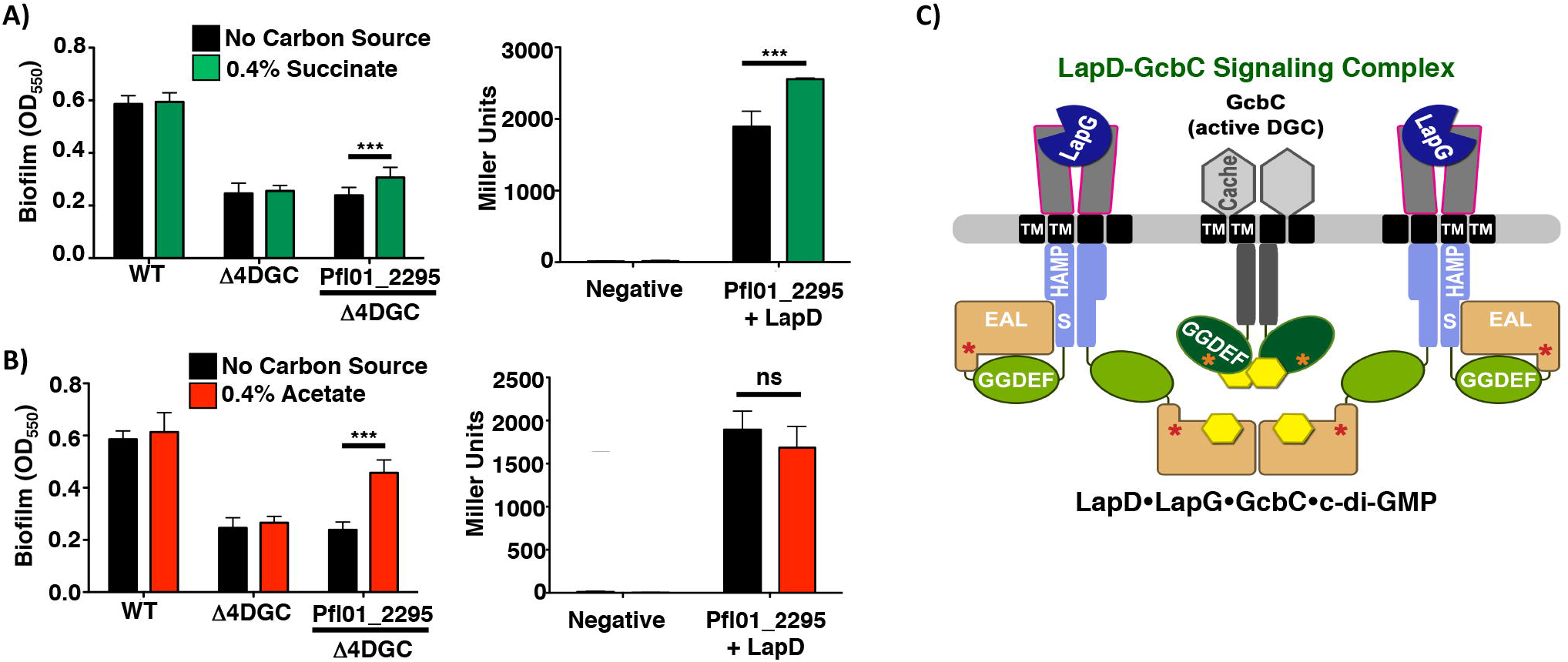
Identification of potential ligands sensed by Pfl01_2295. Shown are the results of biofilm assays and B2H assays with the indicated strains and proteins. For each panel, the presence and absence of each organic acid was compared for both biofilm assay and B2H assay. Panel A = +/-succinate; B = +/- acetate. Biofilm assay data are from six biological replicates (+ SD). B2H assays were performed in triplicate (+ SD). Linear models implemented in R (36) were used to identify organic acid supplemented media whose properties significantly differed from the base medium for both biofilm assay (K10T-1 minimal medium, see Materials and Methods) and B2H assay experiments. P values of <0.05 were considered significant. P<0.01 (**); P<0.001 (***). (C) A model of a GcbC, LapD and LapG “basket” signaling complex (25). In this model, the DGC- and PDE-like domains are indicated by the GGDEF and EAL residues typically associated with the active enzymes; we showed previously that these domains of LapD are not active. Instead, the PDE-like domain of LapD binds c-di-GMP (yellow hexagon). The GGDEF domain of GcbC functions as an active DGC in this context. Also indicated are the HAMP signal transduction domain and the S helix of LapD, the CACHE domain of GcbC, the transmembrane (TM) domains, and I-sites (red and orange stars). The LapG pepriplasmic protease is also shown.

### Discussion

A key open question relating to c-di-GMP signaling is how specificity of a particular output can be mediated in the context of up to dozens of enzymes or receptors making, breaking and binding this dinucleotide second messenger. Here, our data suggest two related mechanisms for controlling output specificity. First, GcbC shows little activity unless it is coexpressed with its receptor. Furthermore, we show that an extracellular ligand can modulate the activity of a GcbC and does so via promoting interaction of the DGC with its cognate receptor. Our data are consistent with the model that citrate or isocitrate binding to the CACHE domain of GcbC enhances interaction with LapD. We propose the increased interaction between GcbC and LapD to have two important consequences: stabilization of a complex that allows direct transfer of the GcbC-generated c-di-GMP signal to the LapD receptor, and equally importantly, activation of the DGC activity of GcbC. Our data suggest that for at least this DGC, GcbC must be in complex with its receptor to be activated. An additional boost in cyclic nucleotide production was measured when citrate or isocitrate was added to the medium in a strain co-expressing GcbC and LapD, indicating that the enhanced GcbC-LapD interaction also enhanced DGC activity. Furthermore, recent work from the Sondermann lab (25) identified a LapD dimer-of-dimers as a c-di-GMP- and LapG-bound state, which could serve as an interaction platform or “receptor basket” for GcbC, thereby forming a large LapD-LapG-GcbC-c-di-GMP signaling complex (Fig. 5C). Our data adds to this model a ligand-based mechanism to enhance the interaction between GcbC and LapD, and thus signaling specificity in the context of a large signaling network.

We suggest a possible mechanism whereby c-di-GMP production by GcbC is promoted when this DGC is in complex with LapD. Our data here suggest that as a baseline condition, GcbC activity is low, a finding consistent with a previous study in which low levels of c-di-GMP synthesis by GcbC were detected (21). Surprisingly, we detected little c-di-GMP production by two different l-site-proximal variants of GcbC when this enzyme was expressed on its own, consistent with the idea that this enzyme is likely inactive when not engaging its receptor. How then does the GcbC-LapD interaction stimulate GcbC activity? We reported previously the somewhat unexpected finding that the I-site of GcbC is required for full LapD-GcbC interaction (13). Based in part on this finding, we propose that the I-site of this diguanylate cyclase (Fig. 5C, orange star in GcbC) may engage LapD (rather than c-di-GMP) to help promote stabilization of the active form of the GcbC. In this model, we propose that it is only when local c-di-GMP levels become very high that GcbC is driven towards the inactive state via I-site binding by the dinucleotide. The addition of citrate may further stabilize the active conformation of GcbC, both by enhancing GcbC-LapD interaction via the GcbC I-site (and likely other interactions (13)) and promoting higher levels of c-di-GMP production.

We have identified two C6, three carboxyl group-containing organic acids, citrate and isocitrate, as potential ligands that bind to GcbC via its CACHE domain. There are an abundance of CACHE domains present as components of signal transduction proteins (16), including proteins involved in c-di-GMP signaling, and yet, the role of CACHE domains in regulating DGC activity is poorly understood. Our data show that citrate and isocitrate are likely ligands for the CACHE domain associated with GcbC. *P. fluorescens*, a plant-growth promoting microbe, forms biofilms on tomato roots; root exudates contain a high abundance of low molecular weight organic acids, including citrate (26, 27, 28). We suggest that many compounds found in root exudates, including additional organic acids, sugars, and metals, may be used as signals by *P. fluorescens* to promote biofilm formation.

As we have shown here, GcbC appears to sense citrate and isocitrate, but not a variety of other structurally related molecules. It is possible that the CACHE domain of GcbC responds to additional signals. For example, in *P. syringe* pv. *actinidiae*, the CACHE domain PscD (PDB ID: 5G4Z) binds glycolate, acetate, propionate, and pyruvate (22), indicating that CACHE domains allow for some promiscuity in their binding of ligands. GcbC has been shown to be part of a large signaling network, interacting with a phosphodiesterase and multiple dual domain containing proteins (20), thus also opening the possibility that different ligands sensed via the CACHE domain of GcbC could dictate which proteins interact with this DGC.

We showed that while GcbC responds to exogenous citrate and isocitrate, other CACHE domain-containing DGCs do not respond to these organic acids. We identified the RXYF motif as important for signal sensing by GcbC. Furthermore, the residue R162, but not R172, is conserved among the CACHE domains of GcbC, Pfl01_2295, Pfl01_2297, and Pfl01_3800 (Fig. S6), thus this variation in sequence likely speaks to the inability of these other DGCs to respond to the same ligands as GcbC. Consistent with the idea that these other CACHE domains bind other signals, we have identified acetate and succinate as potential ligands that are sensed via the CACHE domain of Pfl01_2295. Only succinate enhanced Pfl01_2295-LapD interaction, while acetate promoted Pfl01_2295-dependent biofilm formation but did not enhance LapD interaction. Thus, ligand binding may exert a variety of effects on target proteins. Such a finding may not be surprising in the context of this complex c-di-GMP network, as multiple pathways contribute to biofilm formation and many DGCs have multiple interaction partners (20). Additional work is required to identify all of the ligands that impact the c-di-GMP network in this organism.

Together, our data indicate that extracellular ligands, via their ability to impact protein-protein interactions and/or DGC activity, can modulate c-di-GMP signaling specificity. We think it is unlikely that all nine DGCs shown to interact with LapD do so simultaneously and with high affinity. Rather, we envision a scenario wherein a “cloud” of DGCs with low activity perhaps weakly interact with their receptor(s); a specific DGC-receptor interaction may increase in response to appropriate environmental signals, concomitantly boosting c-di-GMP production, ligand-specific signaling and biofilm formation. Overall, our work provides insight into a ligand-mediated mechanism conferring signaling specificity within a complex network of enzymes and receptors that make, break, and bind c-di-GMP. Given the large number of ligand-binding domains associated with c-di-GMP-metabolizing proteins, the data presented here could represent a general means of regulating c-di-GMP-controlled outputs by enhancing specific interactions between c-di-GMP-metabolizing enzymes and their effectors.

## Material and Methods

### Strains and Media

Bacterial strains used in this study are listed in Table S2, and were cultured and maintained in lysogeny broth (LB) or on 1.5% agar LB plates. *P. fluorescens* was grown at 30°C and *P. aeruginosa* and *E. coli* was grown at 37°C. *E. coli* S17-1-λ-pir was used for maintenance and transfer of plasmids. *Saccharomyces cerevisiae* strain InvSc1 was used for plasmid modification as described previously (29, 30). K10T-1 medium was prepared as described previously (31). Sodium citrate was added to 1.5% agar LB plates and K10T-1 media to a final concentration of 13.6 mM (0.4% wt/vol) for all the experiments described. All organic acids were set at 0.4% wt/ vol and the following concentrations were set as indicated: sodium acetate (29.4mM), pyruvic acid (45.4mM), sodium succinate (14.8mM), sodium fumarate (25.0mM), α-ketoglutarate (27.4mM), and isocitrate (15.5mM). The following antibiotics were used as indicated: gentamycin (15μg/mL for *E. coli*, 30μg/mL for *P. fluorescens* and *P. aeruginosa)*, kanamycin (50μg/mL for *E. coli)*, and carbenicillin (50μg/mL for *E. coli).*

### Biofilm Assay

Biofilm assays were performed as described previously (1). *P. fluorescens* Pf0-1 strains were incubated in K10T-1 minimal medium with and without 0.4% organic acid, as indicated, for 6 hours at 30°C. Biofilms were stained with 0.1% crystal violet, washed with water and then solubilized with a 45% methanol, 45% dH_2_0, and 10% glacial acetic acid solution. The optical density (OD) of the solubilized crystal violet solution was measured at 550 nm to determine the amount of biofilm formed.

### Dot Blot for LapA Localization Assay

Localization of LapA to the cells surface was measured using a HA-tagged LapA variant integrated into the chromosome of *P. fluorescens* as described previously with slight modification (7,10, 32). Bacterial cultures were grown overnight in LB and then subcultured into 5mL K10T-1 at a 1:50 dilution for 6 hours at 30°C. To test how sodium citrate effected LapA localization to the cell surface, sodium citrate was added at the beginning of the 6-hour subculturing period. After 6 hours of incubation, cells were normalized to the lowest OD value, washed twice in K10T-1, and 5μl aliquots were spotted onto a nitrocellulose membrane. Once dried, HA-tagged LapA was probed for by Western blot analysis.

### Bacterial Two-Hybrid Assay

Bacterial two-hybrid (B2H) assays were performed using *E. coli* BTH101 cells based on a previously described system (33). Briefly, ~100 ng of each bacterial two hybrid plasmid was cotransformed into *E. coli* BTH101 by electroporation. *E. coli* BTH101 cells were incubated on LB agar supplemented with 50μg/mL kanamycin, 50μg/mL carbenicillin, and 0.5mM isopropyl β-D-1-thiogalactopyranoside (IPTG) for 24 hours at 30°C. At 24 hours, either β-galactosidase or c-di-GMP levels were quantified as described below, β-galactosidase assays were performed as exactly described previously (12) to quantify the extent of protein-protein interaction, β-galactosidase levels are presented in Miller Units.

### c-di-GMP Quantification Assay

c-di-GMP was extracted from *E. coli* BTH101 cells after incubation on LB agar plates at 30°C for 24 hours. The cells were scraped from the plate surface with 1 mL of dH_2_O, then pelleted and resuspended in 0.250 mL nucleotide extraction buffer (40% methanol, 40% acetonitrile, 20% dH_2_O, and 0.1N formic acid), followed by incubation at - 20°C for 1 hour. Cells were pelleted again and the reaction was neutralized by transfer of 0.2 ml nucleotide extract to 8μl of 15% NH_4_CO_3_. Nucleotide extracts were vacuum-dried and resuspended in 0.2 mL HPLC grade H_2_O. c-di-GMP concentration was analyzed by liquid chromatography-mass spectrometry and compared to a standard curve of known c-di-GMP concentration, as reported (12). The mols of c-di-GMP were normalized to the dry weight of the cell pellet from which the nucleotides were extracted.

## Congo Red Binding Assay

Assessment of DGC activity using the congo red binding assays was performed as described previously (12).

## Swim Motility Assay

Swim motility assays were performed as described previously (21). K10T-1 plates containing 0. 35% agar were prepared by adding filter-sterilized K10T-1 media to an autoclaved, molten agar solution and plates were solidified for 3 hours. Overnight cultures were normalized to the lowest OD value and washed twice in K10T-1 medium. A 2-200μl pipette micropipette tip was used to inoculate each plate. A sterile pipette tip was dipped into each strain and plunged halfway into the swim agar plate. Plates were incubated at 30°C for ~30 hours and plates were photographed. The swim area was calculated using ImageJ software.

## Growth Curve Assay

The amount of growth of *P. fluorescens* in the presence and absence of citrate was measured. Bacterial cultures were grown overnight in liquid LB and then subcultured in 5mL K10T-1 with and without 0.4% citrate at a 1:50 ratio for 6 hours at 30°C. Every two hours, the OD of the bacterial culture was measured at 600nm to determine the amount of growth that occurred.

## Acknowledgement

We thank members of the lab for helpful discussions, the reviewers for their helpful comments, and Tom Hampton for assistance with statistical analysis. This work was supported by the NIH via grant R01GM123609 (to H.S. and G.A.O.) and P20 RR030360.

## Supplemental Material

**Fig. SI. CLUSTAL alignment of the amino acid sequence of the CACHE domain of GcbC and rpHK1S-Z16 (PDB ID: 3LIF) of *R. palustris***. The CACHE domain of GcbC and rpHK1S-Z16 showed 31% identity at the amino acid level (E-score = le^−27^). The red amino acids indicate the conserved residues in the proposed citrate-binding pocket. A subset of these residues are mutated (see Figures 2 and 3 in the main text, and Table S1).

**Fig. S2. Effect of a *lapA* mutation on citrate-mediated biofilm formation.** (A) Quantitative analysis of dose-dependent biofilm formation by WT *P. fluorescens* by addition of increasing concentrations of citrate. Citrate concentrations of 0%, 0.4%, and 0.8% were tested. (B) Quantitative analysis of biofilm formation by the WT and *lapA* mutant of *P. fluorescens* in the presence and absence of citrate. (C) Quantification of LapA levels in whole cell lysate in the presence and absence of 0.4% citrate by WT *P. fluorescens* Pf0-1. Representative blots are shown. Data shown is the average of three replicates (+ SD). (D) Growth of WT *P. fluorescens* in the presence and absence of citrate for 6 hours at 30°C. (E) Motility by WT *P. fluorescens* in the presence and absence of citrate at 30°C for 30 hours using a soft agar (0.3%) motility assay. Representative images are shown. In all experiments in this figure, assays were performed in triplicate (+ SD), and horizontal black bar indicates a P value of <0.05 (*), <0.01 (**), or <0.001 (***) by a student’s t-test comparing each strain without citrate versus with added citrate, or in panel A, comparing the two different citrate concentrations tested.

**Fig. S3. Assessment of DGC activity via congo red binding.** Qualitative analysis of GcbC, Pfl01_2295, and Pfl01_2297 to test for functional c-di-GMP production as described in the main text. These plasmids were transformed into *P. aeruginosa* PAM, and the strains grown on congo red plates supplemented with 0.1% arabinose at 37°C to induce expression of the plasmid-borne DGCs for 24 hours. The binding of red pigment indicates Pel exopolysaccharide production, which serves as an indirect means to assess the amount of c-di-GMP produced by the indicated strain. GcbC and Pfl01_2295 show a higher degree of congo red binding as compared to the empty pMQ72 control and the *pelA* deletion mutant strain. Pfl01_2297 showed only a very modest enhancement of congo red binding compared to the empty pMQ72 control and *pelA* deletion mutant strain.

**Fig. S4. Modeling the CACHE domain of GcbC on rpHKlS-Zlθ (PDB ID: 3LIF) and vpHK1S-Z8 (PDB ID: 3LID).** Shown is the model of the ligand-binding site of rpHK1S-Z16 (PDB ID: 3LIF) and vpHK1S-Z8 (PDB ID: 3LID), as well as the homology model of the ligand-binding site of the apo-GcbC. On the far right is shown the overall model of the GcbC periplasmic domain based on the structure of rpHK1S-Z16 (PDB ID: 3LIF).

**Fig. S5. Identification of ligands sensed by GcbC’s CACHE domain.** (A) Shown is a list of organic acids that are color coordinated with the graph in panel B. The chemical structure of acetate, pyruvate, succinate, fumarate, α-ketoglutarate, isocitrate, and citrate are shown. The number of carbon atoms of each compound is listed on the right. (B) Quantitative analysis of biofilm formation by WT *P. fluorescens* and the Δ4DGC mutant strain in the presence of 0.4% acetate, pyruvate, succinate, fumarate, α-ketoglutarate, and isocitrate. The empty vector control and a plasmid carrying GcbC were introduced into the Δ4DGC mutant strain. Biofilm assay data are from six biological replicates (+ SD). Linear models implemented in R (36) were used to identify organic acid supplemented media whose properties significantly differed from the base medium (K10T-1 minimal medium, see Materials and Methods) for both biofilm assay and B2H assay experiments. The reduced biofilm levels for medium supplemented with α-ketoglutarate is due to the poor growth of the strains in the presence of this compound (not shown). P values of <0.05 were considered significant. P<0.05 (*); P<0.001 (***). (C) Biofilm formation by the indicated strains + isocitrate. The Δ4DGC mutant strain is used with WT GcbC and GcbC-R139E variant introduced on plasmids. For panel C, experiments were performed in triplicate (+ SD), and horizontal black bars indicate a P value of <0.05(*) with a student’s t-test comparing the presence and absence of isocitrate amongst each strain.

**Fig. S6. CLUSTAL alignment of the amino acid sequence of the CACHE domains of GcbC, Pfl01_2295, and Pfl01_2297.** The red amino acids indicate the conserved residues of the RXYF motif. The residues highlighted in blue are proposed to make up the putative citrate-binding pocket of the CACHE domain of GcbC. The RXYF motif and residues R139 and R162 of GcbC are conserved in the CACHE domains of Pfl01_2295, Pfl01_2297, and Pfl01_3800.

**Fig. S7. Identification of potential ligands sensed by Pfl01_2295.** Shown are the results of biofilm assays with the indicated strains. For each panel, the presence and absence of each organic acid was compared for each biofilm assay. Panel A = +/-isocitrate; B = +/-pyruvate; C = +/-fumarate. Biofilm assays are representative of six biological replicates P<0.001 (***).

**Table SI. CACHE Domain Mutations. Table S2. Strains, Plasmids, and Oligonucleotides.**

